# Core gastric microbiota linked to pathogenesis and preserved across age stratified cohorts

**DOI:** 10.1101/2025.08.26.672344

**Authors:** Weidong Huang, Qiujing Chen, Xuejie Gao, Yongyuan Su, Jiayi Shen, Tao Zhang, Hao Zhang, Yanbin Chen, Yingying Yu, Wen Wang, Hewei Jiang, Xu Lin, Zhaowei Xu

**Author notes:** Corresponding authors. (Zhaowei Xu), (Xu Lin), (Hewei Jiang). These authors contributed equally to this study (Weidong Huang, Qiujing Chen, Xuejie Gao).

## Abstract

Mounting evidences reveal a complex microbial niche in gastric environment. Nevertheless, the phylogenetic stability of gastric microbiota remains undetermined. Here, using full-length 16S rRNA and ITS amplicon sequencing applied to gastric brushes from a pediatric endoscopic cohort (n=243), we identified a core microbiota comprising 11 genera, which clustered into two antagonistic communities (CST1A and CST1B). CST1A was enriched in healthy subjects, while CST1B correlated with gastroduodenal disease. Notably, this bipartite architecture arose through bacterial interactions, with fungal taxa demonstrating minimal integration. Further stratified analyses revealed that *H. pylori* infection preferentially depleted CST1A taxa. The subclusters exhibited divergent metabolic profiles, with CST1A specializing in motility processes and CST1B in nucleotide/peptidoglycan synthesis. Cross-cohort validation of six datasets (n=1,183) confirmed the persistent relevance of the CST1A-CST1B structure across disease stages. Collectively, our findings define a core gastric microbiota established in early life, linked to gastric disease risk, and persisting into adulthood.

**Highlights:**

1. Identification of a gastric bacterial network established in early life.
2. Dysbiosis in two subclusters within the core microbiota is associated with pediatric peptic ulcers.
3. *Helicobacter pylori* infection reduces the abundance of the core microbiota, with a particularly greater inhibitory effect on CST1A.
4. The core microbiota is preserved across age stratified cohorts and is associated with gastric cancer.

## Introduction

The human stomach was conceptualized as a physiologically hostile microenvironment for microbial persistence, characterized by its extreme acidic luminal pH (1.5-3.5) and proteolytic activity mediated through pepsinogen activation and gastric lipase^1^. With the advancement of high-throughput sequencing technologies, the stomach was redefined as a dynamic microbial habitat exhibiting stratified ecological organization^2–7^. Notably, *Helicobacter pylori* (*H. pylori*, HP) remains the prototypical gastric pathogen^8^ and emerging studies have identified *Streptococcus anginosus* as a new gastric pathobiont capable of establishing persistent colonization with carcinogenic effects^9^, suggesting a complicated microbial interaction in gastropathology. However, the functional interplay between commensal microbiota remains unclear.

As the principal etiopathogenic driver of gastric adenocarcinoma and peptic ulcer disease (PUD), *H. pylori* can induce chronic inflammation and genomic instability in gastric epithelia^10^. Paradoxically, while global *H. pylori* seroprevalence reaches 43.1%^11^, only 1-3% of colonized individuals progress to gastric malignancy, underscoring the limited contribution of *H. pylori* alone. Intriguingly, longitudinal cohort studies reveal persistent 2.8-fold higher risk gastric cancer risk in post-eradicated patients versus uninfected controls^12^. Furthermore, growing evidence supports shifting the focus from single pathogens to theories of microbial imbalance in the intestine^13,14^. Whether gastric diseases likewise arise from disruption of a microbiota network, rather than from single invaders alone, remains unclear.

Despite recent progress in gastric microbiome research, current microbial profiling methods have limited ability to capture early developmental or physiologically normal states of the gastric microbiota, as these states are often subject to confounding by factors such as advanced age, polypharmacy, and comorbid metabolic disorders^15^. Notably, the development of precancerous lesions induces profound histopathological alterations, including glandular atrophy^16^ and chronic inflammatory infiltration^17^. These alterations create a microenvironment that fundamentally reshapes microbial niches, thereby complicating causal inference in microbiome disease associations^18^. These methodological challenges highlight the critical need for adopting a developmental life course framework. Specifically, prioritizing investigations into early-life microbiome dynamics could reveal evolutionarily preserved core taxa that constitute the foundational gastric ecosystem.

Pediatric populations serve as a strategic model system for identifying autochthonous gastric microbiota^19^. The constrained environmental heterogeneity, limited cumulative antibiotic exposure, and homogeneous dietary patterns inherent to childhood development collectively attenuate ecological confounding variables. Furthermore, the pediatric phase represents a critical ontogenetic window characterized by bidirectional host-microbe coevolution^20^, during which microbial communities are more likely to retain ancestral ecological signatures through stable vertical transmission^21^. This synchronicity between host mucosal maturation and microbial colonization suggests that childhood microbiota may serve as phylogenetically informative markers of gastric ecosystem assembly.

In this study, we performed full-length 16S rRNA and ITS amplicon sequencing on gastric brushing specimens from 243 children undergoing upper endoscopy, and systematically delineated the structure of the core gastric microbiota and its links to disease. Our analyses revealed a preserved microbiota network that emerges in early life and persists into adulthood. Within this network, two functionally antagonistic subclusters form a “seesaw” configuration whose dysbiosis is tightly coupled to pediatric peptic ulcer pathology. *H. pylori* colonization suppresses the core gastric microbiota, with persistent disruption of this ecosystem propagating across the gastric cancer continuum. Together, these discoveries establish a life course ecological framework for gastric microbial stability, pinpoint specific network elements that forecast distinct disease trajectories.

## Results

### Identification of gastric microbial network in *H. pylori*-negative pediatric

A total of 246 children who underwent upper gastrointestinal endoscopy were enrolled in this study, including 154 males and 92 females. Among them, 167 were delivered vaginally and 77 via cesarean section. Based on endoscopic diagnoses, 187 were identified with superficial gastritis and 59 with duodenal ulcer **(Table 1)**. To assess *H. pylori* infection status, we performed high-throughput sequencing and compared the results with conventional diagnostic methods, including the ¹³C-urea breath test, immunohistochemistry, and rapid urease test. Receiver operating characteristic (ROC) curve analysis yielded an optimal read-based cutoff value of 0.01, achieving an area under the curve (AUC) of 0.927 (95% CI: 0.855-1.000) **(Fig. S1A)**. Based on this threshold and endoscopic findings, participants were classified into four groups: Hp-negative non-ulcer controls (HpNNC, n = 149), Hp-negative duodenal ulcer (HpNDU, n = 29), Hp-positive non-ulcer controls (HpPNC, n = 38), and Hp-positive duodenal ulcer (HpPDU, n = 28). Given the hematological markers for duodenal ulcer, we compared blood parameters between ulcer and non-ulcer groups, adjusting for age, sex, and delivery mode. Children in the control group exhibited significantly higher mean corpuscular hemoglobin concentration (MCHC), hematocrit (HCT), red blood cell count (RBC), and hemoglobin (HGB), while those in the ulcer group showed elevated immature granulocyte (IG) counts, red cell distribution width (RDW), and RDW coefficient of variation (RDW-CV) **(Fig. S1B)**. These clinical distinctions validated the disease stratification and provided phenotypic references for microbial correlation analyses.

**Table 1.**
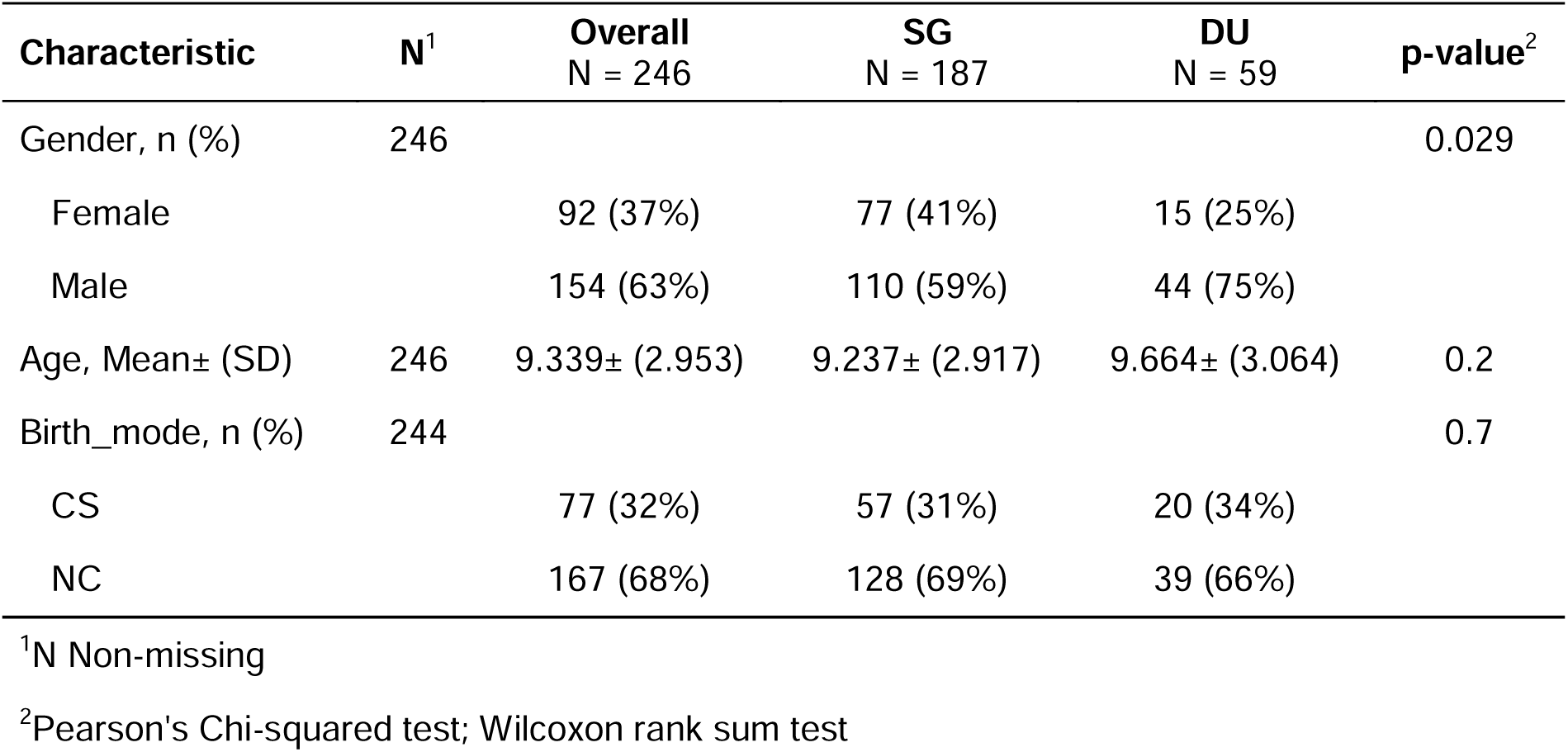
Baseline characteristics and endoscopic diagnoses of the study population. Data are presented as n (%). The study included 246 children aged between 2-17 years, with 154 males (62.6%) and 92 females (37.4%). The mode of delivery included 167 vaginal births (67.9%) and 77 cesarean sections (31.3%). Endoscopic diagnoses based on gastroscopy revealed 187 cases of superficial gastritis (76.0%) and 59 cases of duodenal ulcer (24.0%).

| **Characteristic** | **N<sup>1</sup>** | **Overall**<br>**N = 246** | **SG**<br>**N = 187** | **DU**<br>**N = 59** | **p-value<sup>2</sup>** |
| --- | --- | --- | --- | --- | --- |
| Gender, n (%) | 246 |  |  |  | 0.029 |
| Female |  | 92 (37%) | 77 (41%) | 15 (25%) |  |
| Male |  | 154 (63%) | 110 (59%) | 44 (75%) |  |
| Age, Mean± (SD) | 246 | 9.339± (2.953) | 9.237± (2.917) | 9.664± (3.064) | 0.2 |
| Birth_mode, n (%) | 244 |  |  |  | 0.7 |
| CS |  | 77 (32%) | 57 (31%) | 20 (34%) |  |
| NC |  | 167 (68%) | 128 (69%) | 39 (66%) |  |

<sup>1</sup>N Non-missing
<sup>2</sup>Pearson's Chi-squared test; Wilcoxon rank sum test

To identify the core gastric microbiota, we employed the full-length 16S rRNA and ITS amplicon sequencing on samples from a pediatric endoscopic cohort (n=243) through third-generation long read platform. The microbial abundance matrices were denoised by removing genera with relative abundance <0.005 and retaining only those present in ≥10% of samples **(Fig. S1C)**. Genus-level co-occurrence networks were independently constructed for HpNNC and HpNDU groups using Spearman correlations (ρ > 0.5, *q* < 0.05). The HpNNC network comprised 24 nodes and 83 edges, while the HpNDU network expanded to 27 nodes and 133 edges **(Fig. 1B)**. Across both networks, 67 stable genus-genus correlations (45 positive, 22 negative) were identified, involving 23 genera **(Fig. 1C)**. Hierarchical clustering of these genera revealed two major clusters, CST1 and CST2. Notably, CST1 comprised 21 genera and could be further divided into two opposing submodules: CST1A and CST1B, characterized by strong positive intra-submodule correlations and negative inter-submodule associations **(Fig. 1D)**. To validate the clinical relevance of this network, multivariate regression analyses (adjusted for age, sex, and delivery mode) were conducted. Only the CST1 cluster showed significant correlations with host hematological parameters: CST1A was positively associated with both HGB and HCT (*q* < 0.001), whereas CST1B exhibited negative associations. No significant correlations were observed for CST2 **(Fig. 1E)**. These results collectively support the robustness and host relevance of the CST1 microbial cluster.

**Figure 1.**
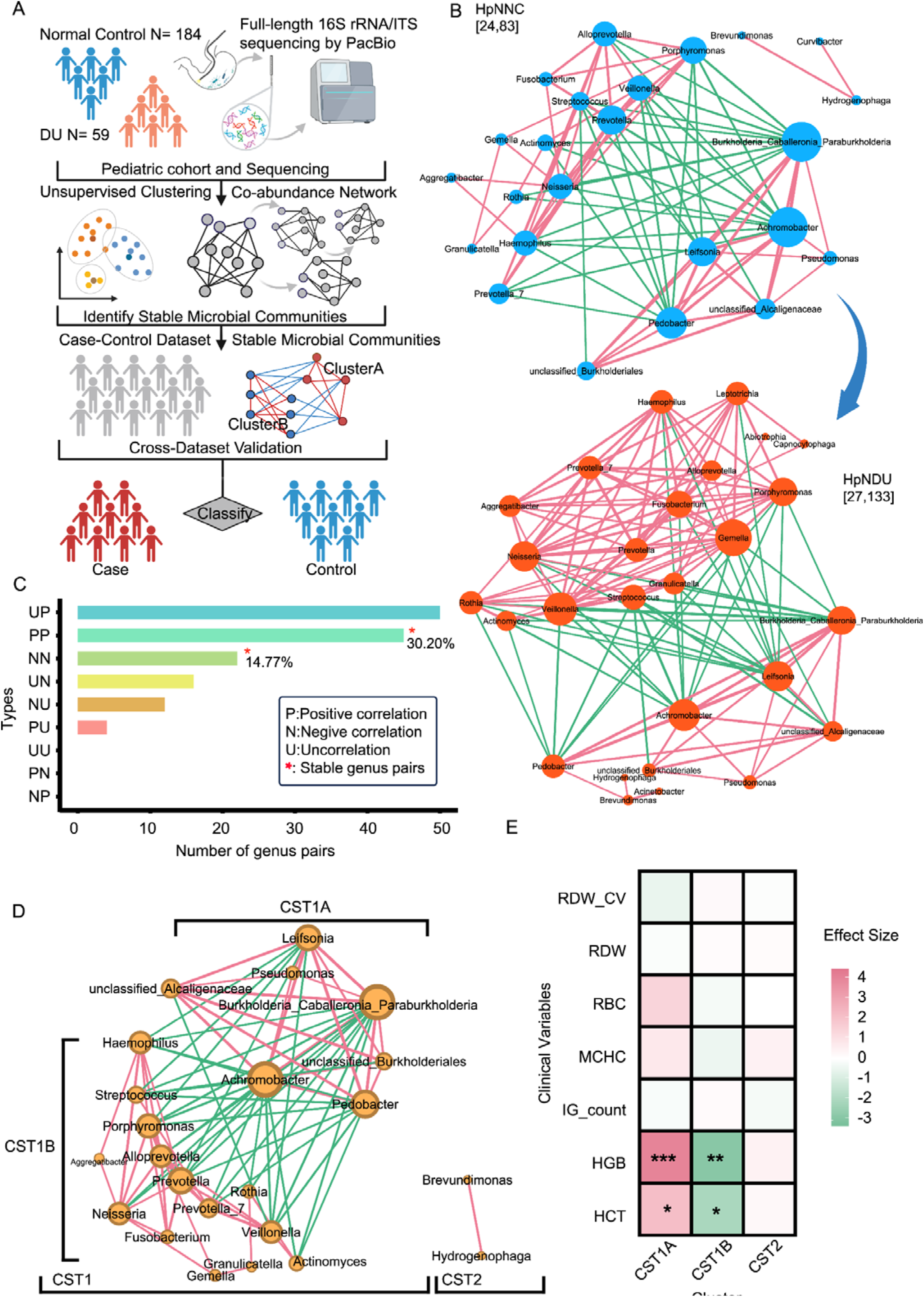
Identification of gastric core microbial genera and modular structure in *H. pylori*-negative children. (A) Overview of the study design: full-length 16S rRNA/ITS amplicon sequencing was performed on 246 pediatric gastric mucosal samples to profile bacterial and fungal communities. (B) Genus-level co-occurrence networks were independently constructed in *H. pylori*-negative normal controls (HpNNC) and duodenal ulcer patients (HpNDU) using Spearman correlation (r > 0.5, *q* < 0.05). The HpNNC network contained 24 nodes and 83 edges; The HpNDU contained 27 nodes and 133 edges. (C) A total of 67 shared genus-pairs (45 positive, 22 negative) were identified between the two networks, involving 23 genera. Stable correlations across disease stage point (NNN and PPP), were indicated by asterisks. (D) Hierarchical clustering of the 23 genera revealed two major ecological clusters: C1 and C2. Cluster C1 further split into two negatively correlated submodules: CST1A and CST1B. Intra-module correlations were positive; inter-module correlations were predominantly negative. (E) Multivariable linear regression (adjusted for age, sex, and birth mode) indicated that only Cluster C1 showed significant associations with clinical hematological indices: CST1A was positively correlated with HGB and HCT, whereas CST1B was negatively correlated. No significant associations were observed for Cluster C2. Microbiome-phenotype associations were analyzed using MaAsLin2 (v1.16.0) with a linear mixed-effects model (fixed effects: disease status, age, gender, and birth mode). Microbial abundance data were normalized via centered log-ratio (CLR) transformation. Significance thresholds were defined as FDR-adjusted *q* < 0.05 using Benjamini-Hochberg correction. Significance levels in figures are denoted as: * *q* < 0.05, \*\**q* < 0.01, \*\*\**q* < 0.001. (Benjamini-Hochberg correction).

### Cross validation of the core gastric microbiota in *H. pylori*-infected pediatric ulcer

To assess whether the preserved core microbial network identified in *H. pylori*-negative children remained stable under *H. pylori* infection, we constructed genus-level co-occurrence networks for the HpPNC and HpPDU groups. The HpPNC network comprised 20 nodes and 49 edges, whereas the HpPDU network expanded to 33 nodes and 144 edges. Notably, *Helicobacter* exhibited distinct antagonistic patterns across disease states: in HpPNC, it showed negative correlations primarily with *Streptococcus*-dominated taxa, while in HpPDU, its antagonism extended to a *Burkholderia-Caballeronia-Paraburkholderia (BCP)*-dominated group **(Fig. 2A)**. Tracking 15 genera across the HpNNC-HpPNC-HpPDU continuum revealed 26 stable positive correlations **(Fig. 2B)**, all of which belonged to the previously defined 23 genera from the *H. pylori*-negative core network **(Fig. 2C)**. These results indicate that, despite the presence of *H. pylori*, a subset of core bacterial interactions remains preserved, albeit with altered antagonistic targets.

**Figure 2.**
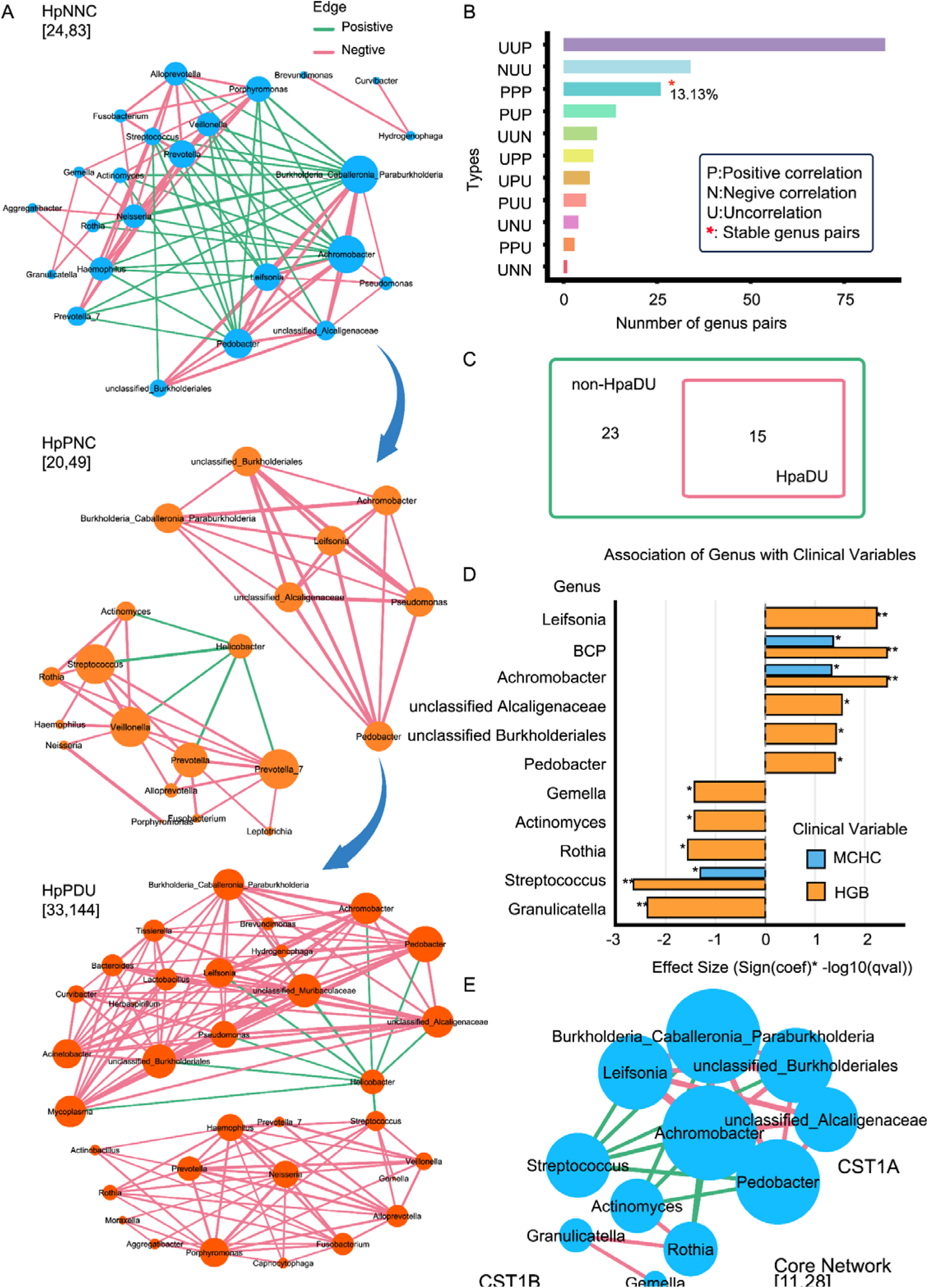
Cross-validation of the core network structure in *H. pylori*-positive subgroups and identification of 11 core genera. (A) Genus-level co-occurrence networks constructed in Hp-positive normal (HpPNC) and duodenal ulcer (HpPDU) groups. The HpPNC network contained 20 nodes and 49 edges; HpPDU contained 33 nodes and 144 edges. *H. pylori* exhibited negative correlations with *Streptococcus* in HpPNC and with *Burkholderia Caballeronia Paraburkholderia* (*BCP*) in HpPDU. (B) Across HpNNC-HpPNC-HpPDU stages, 26 genus pairs remained positively correlated, involving 15 genera. (C) These 15 genera overlapped with the 23 genera identified in *H. pylori*-negative networks, supporting structural stability. Stable correlations across dieases stage points (NNN and PPP), were indicated by asterisks. (D) Multivariable analysis (adjusted for age, sex, and birth mode) confirmed significant associations (*q* < 0.05) between 11 genera and hematological indicators. Six genera (*Leifsonia*, *Burkholderia Caballeronia Paraburkholderia*(*BCP*), *Achromobacter*, *unclassified Alcaligenaceae*, *unclassified Burkholderiales*, and *Pedobacter*) were assigned to CST1A (positively correlated with HGB; *Burkholderia Caballeronia Paraburkholderia*(*BCP*) and *Achromobacter* also with MCHC), whereas five (*Gemella*, *Actinomyces*, *Rothia*, *Streptococcus*, and *Granulicatella*) were assigned to CST1B (negatively correlated with HGB; *Streptococcus* also with MCHC). Microbiome-phenotype associations were analyzed using MaAsLin2 (v1.16.0) with a linear mixed-effects model (fixed effects: disease status, age, gender, and birth mode). Microbial abundance data were normalized via centered log-ratio (CLR) transformation. Significance thresholds were defined as FDR-adjusted q < 0.05 using Benjamini-Hochberg correction. Significance levels in figures are denoted as: \**q* < 0.05, ** *q* < 0.01, \*\*\**q* < 0.001. (Benjamini-Hochberg correction). (E) Core network reconstructed using the 11 core genera, showing positive intra-module and negative inter-module correlations between CST1A and CST1B.

To further delineate the clinical relevance of the core taxa, we examined associations between the 23 genera and hematological parameters that differed significantly between DU and control groups. Multivariate linear regression (adjusted for age, sex, and delivery mode) identified 11 genera significantly correlated with at least one clinical parameter (*q* < 0.05), including *Leifsonia*, *BCP*, *Achromobacter*, *unclassified Alcaligenaceae*, *unclassified Burkholderiales*, *Pedobacter*, *Gemella*, *Actinomyces*, *Rothia*, *Streptococcus*, and *Granulicatella*. CST1A genera (e.g., *Leifsonia*, *BCP*, *Achromobacter*) were positively associated with hemoglobin (HGB), with *BCP* and *Achromobacter* also linked to mean corpuscular hemoglobin concentration (MCHC). In contrast, CST1B genera (e.g., *Streptococcus*, *Rothia*) were negatively associated with both HGB and MCHC **(Fig. 2D)**. Using these 11 clinically relevant genera, we reconstructed a refined core microbial network, which preserved the original modular layout: strong positive correlations within each submodule (CST1A and CST1B) and negative correlations between them **(Fig. 2E)**. These refined modules reinforce the clinical relevance of the original CST1 structure and support its robustness as a functionally organized microbial hub.

In addition, we profiled the gastric fungal community within the HpNNC group. Across all samples, the five most abundant fungal genera were *Malassezia*, *Candida*, *Cutaneotrichosporon*, *Aspergillus*, and *Trametes* **(Supplementary Material, page 1)**. After applying abundance (0.006) and prevalence (0.08) thresholds **(Fig. S1D)**, fungal co-occurrence networks were constructed for the HpNDU and HpPDU groups (ρ > 0.2, *q* < 0.05). However, no significant associations were detected in the HpPDU network. Moreover, fungal networks in HpNNC (4 nodes, 2 edges), HpPNC (17 nodes, 6 edges), and HpNDU (10 nodes, 33 edges) displayed no consistent topological features **(Supplementary Material, pages 2-4)**. These findings suggest a lack of stable fungal interaction networks in the gastric environment, in contrast to the structured bacterial communities.

### Modular subtyping reveals microbial distribution among superficial gastritis patients

To explore inter-individual variability in the gastric microbiota, we stratified the HpNNC group defined as *H. pylori*-negative individuals diagnosed with superficial gastritis, based on features derived from the previously defined 11 genus core microbial network. Microbial composition within the HpNNC group exhibited significant clustering patterns **(Fig. 3A)**. Partitioning Around Medoids (PAM) clustering identified two optimal clusters (K = 2), as supported by silhouette width analysis **(Fig. 3B, Fig. S2A)**. Cluster robustness was confirmed using k-means clustering, yielding a high adjusted Rand index (ARI = 0.84; **Fig. S2B**). Based on this, the HpNNC group was divided into two subgroups: HpNNC1 (n = 71) and HpNNC2 (n = 78). We next compared hematological profiles between each HpNNC subtype and the *H. pylori*-negative duodenal ulcer group (HpNDU), adjusting for age, sex, and delivery mode. Although the differences were not statistically significant across all measures, HpNNC1 tended to show greater divergence from HpNDU compared to HpNNC2, with relatively higher hemoglobin (HGB, Estimate = +11.37, *q* < 0.01), increased red blood cell count (RBC, Estimate = +0.29, *q* < 0.05), and reduced red cell distribution width (RDW, Estimate = -2.51, *q* < 0.05) **(Fig. 3C)**. Compositional analysis revealed that HpNNC1 was enriched in CST1A-associated genera, such as *Achromobacter* and *Burkholderia-Caballeronia-Paraburkholderia (BCP)*, whereas HpNNC2 was dominated by CST1B genera, including *Streptococcus* and *Rothia* **(Fig. 3D)**. Based on these taxonomic differences, we designated HpNNC1 as the AB subtype (CST1A-enriched) and HpNNC2 as the S subtype (CST1B-enriched). All signature genera showed >25% prevalence within their respective subtypes **(Fig. 3E)**. These observations suggest that the AB and S microbial subtypes exhibit distinct ecological and clinical trends, though further validation is needed to establish their potential clinical significance.

**Figure 3.**
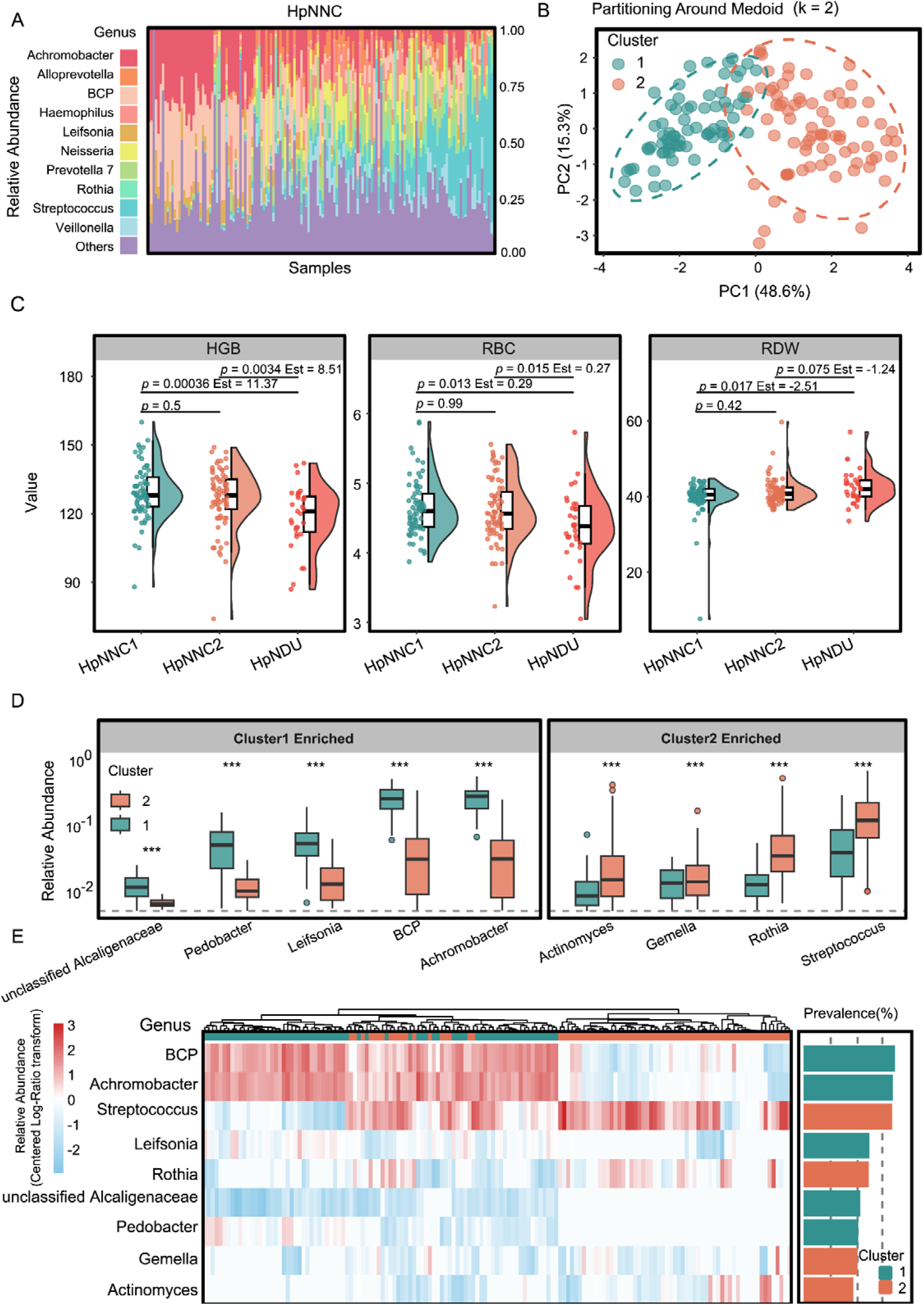
Identification of two microbial subtypes in *H. pylori*-negative controls and their clinical relevance. (A) Stacked bar plot showing the relative abundance of the top 10 genera in the *H. pylori*-negative normal control (HpNNC) group. Each bar represents an individual sample; the x-axis denotes sample ID, and colors represent genera. (B) Based on the abundance matrix of 11 core genera (as defined in Fig. 2E), partitioning around medoids (PAM) clustering identified two robust microbial subtypes (K = 2), supported by high concordance using k-means clustering (see Fig. S2B, adjusted Rand index = 0.84). HpNNC individuals were classified into HpNNC1 (n = 71) and HpNNC2 (n = 78). (C) Multivariable linear regression (adjusted for age, sex, and birth mode) showed that HpNNC1 differed more significantly from the HpNDU group in hematological indices than HpNNC2, including higher HGB (Estimate = 11.37, *q* < 0.01), higher RBC (Estimate = 0.29, *q* < 0.05), and lower RDW (Estimate = -2.51, *q* < 0.05). (D) Box plots showing the relative abundance of representative CST1A and CST1B genera in HpNNC1 and HpNNC2 subgroups. Each box represents the interquartile range (IQR), with the median indicated; whiskers extend to 1.5× IQR. Microbial abundance differential analysis was performed using MaAsLin2(v1.6.0) with robust linear models and fixed effects comprising disease status, age, gender, and birth mode. Raw abundances were normalized by sum scaling, with significance thresholds defined as log_2_(fold change)>1 and FDR-adjusted q<0.05. Differential taxa are annotated in figures as: *q<0.05, **q<0.01, *** q<0.001 (Benjamini-Hochberg correction). (E) Heatmap displaying scaled relative abundance (Z-score) of the 11 core genera across HpNNC1 and HpNNC2 individuals (left panel), alongside a bar plot showing genus-level prevalence (>25%) within each subtype (right panel). Color intensities indicate relative abundance across samples.

### Disruption of the CST1A-CST1B network balance in ulceration and its interplay with *H. pylori*

To investigate the disruption of the core gastric microbiota under ulcerative conditions, we used HpNNC1 (CST1A, n = 71) as an optimized reference group and compared microbial profiles across disease states. Members of the CST1A module,such as *Achromobacter* and *Burkholderia-Caballeronia-Paraburkholderia (BCP)* exhibited significantly lower abundance and prevalence in both HpNDU and HpPDU groups compared to HpNNC1 **(Fig. 4A)**. The CST1A/CST1B abundance ratio was also significantly reduced in both HpNDU and HpPDU (p = 0.013), indicating a consistent shift in microbial balance under ulcerative conditions **(Fig. 4B)**. Although no individual genera were significantly different between HpPNC and HpPDU, core genera from the CST1 network still effectively distinguished these groups in classification models (AUC = 0.937, 95% CI: 0.886-0.987; **Fig. S2C-E**). These findings indicate a consistent depletion of CST1A taxa and disruption of the CST1A/CST1B modular balance in ulcer states.

**Figure 4.**
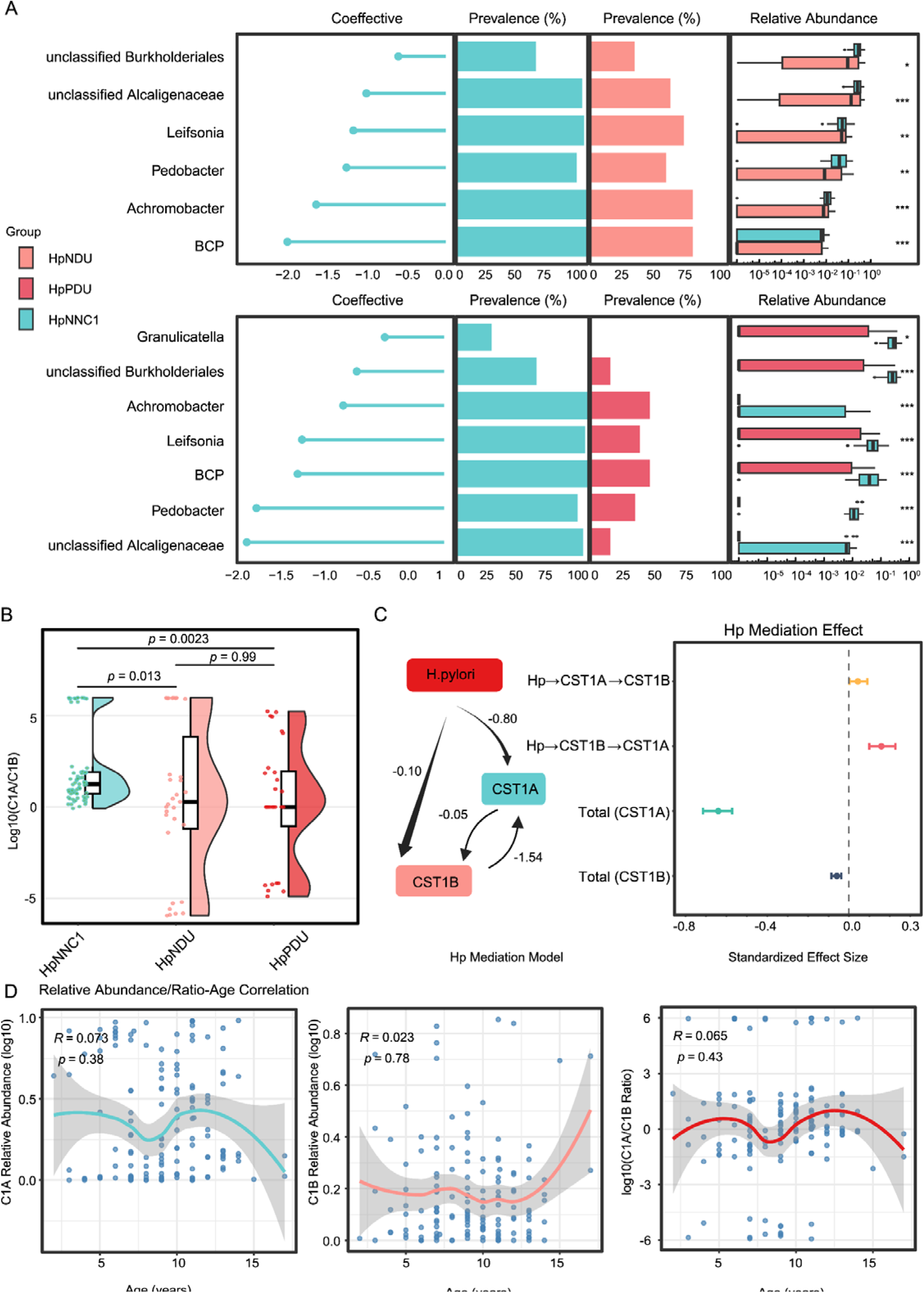
Disruption of the gastric core microbial network in ulcer states and indirect regulation by *H. pylori*. (A) Differential abundance analysis of core genera between the AB-type reference group (HpNNC1) and ulcer groups (HpNDU and HpPDU). The upper and lower panels represent comparisons between HpNNC1 vs. HpNDU and HpNNC1 vs. HpPDU, respectively. Each panel contains four aligned subplots sharing a common y-axis (differential genera): (1) bar plot of model-estimated coefficients (effect sizes); (2) and (3) prevalence of each genus in HpNNC1 and ulcer group, respectively; and (4) box plots showing genus-level relative abundance distributions between groups. Microbial abundance differential analysis was performed using MaAsLin2(v1.6.0) with robust linear models and fixed effects comprising disease status, age, gender, and birth mode. Raw abundances were normalized by sum scaling, with significance thresholds defined as log_2_(fold change)>1 and FDR-adjusted q<0.05. Differential taxa are annotated in figures as: \**q*<0.05, \*\**q*<0.01, *** *q*<0.001 (Benjamini-Hochberg correction). (B) Boxplots showing CST1A/CST1B in ulcer groups compared to HpNNC1. The x-axis represents effect size, and the y-axis indicates -log10-transformed q-values. (C) Panel summarizing bidirectional mediation analysis between *H. pylori* and the core microbial modules (CST1A and CST1B). Left: path diagram of the structural equation model. Arrows represent estimated effect coefficients along each path, including direct effects (Hp → CST1A/CST1B), inter-module suppression (CST1A→CST1B, CST1B→ CST1A), and total indirect effects. Right: bar plot displaying the magnitude and 95% confidence intervals of each direct, indirect, and total effect, highlighting the stronger mediation of *H. pylori* on CST1A via CST1B than the reverse. (D) Correlation scatterplots showing the association between host age and the relative abundance of CST1A, CST1B, and their ratio. Trend lines were fitted using generalized additive models. No significant correlations were observed.

To further examine the regulatory role of *H. pylori*, mediation analysis showed a negative effect on both CST1A (-0.64, 95% CI: -0.707 to -0.574) and CST1B (-0.05, 95% CI: -0.0839 to -0.0375). Bidirectional indirect effects were observed, whereby *H. pylori* influenced CST1B via CST1A (0.0435, 95% CI: 0.0003-0.086), and CST1A via CST1B (0.1587, 95% CI: 0.099-0.22), suggesting multi-pathway modulation of microbial equilibrium **(Fig. 4C)**. To assess the stability of the core microbial modules, we next evaluated associations between CST1A/CST1B abundances and host factors in the *H. pylori*-negative reference group. No statistically significant correlations with age were observed for CST1A abundance (R = 0.073), CST1B abundance (R = 0.023), or the CST1A/CST1B ratio (R = 0.065), with all p-values exceeding 0.05 **(Fig. 4D)**. Similarly, no significant differences were observed by sex or delivery mode (*q* > 0.05; **Supplementary Material, pages 5-6**). Together, these results highlight the CST1A-CST1B dynamic balance in pediatric ulcer and its interplay with *H. pylori*.

### Phylo-functional divergence of CST1A and CST1B reveals a modular gastric ecosystem

To characterize the ecological structure of the core gastric microbiota, we constructed co-occurrence networks spanning genus, species, and OTU levels. The 23 core genera were taxonomically affiliated with four major phyla — *Actinobacteriota*, *Firmicutes*, *Bacteroidota*, and *Proteobacteria*— and exhibited clear differentiation between CST1A and CST1B at the order level **(Fig. 5A)**. Correlation analysis across taxonomic hierarchies (ρ > 0.5, p < 0.05) revealed a structurally preserved pattern, which exhibited positive intra-group correlations but negative inter-group correlations **(Fig. 5B)**. At the species level, CST1A was primarily composed of *Pedobacter nutrimenti* (14%), *Achromobacter spanius*(8%), and unclassified *Burkholderia-Caballeronia-Paraburkholderia* (7%). In contrast, CST1B was dominated by *Rothia mucilaginosa* (14%), *unclassified Streptococcus* (12%), and *Streptococcus salivarius* (8%) **(Fig. 5C)**. Predicted functional profiling suggested distinct metabolic potentials between the two modules: CST1A was associated with cell motility-related functions (*q* = 2.1 × 10□□) and fatty acid degradation pathways (*q* < 0.001), whereas CST1B was linked to amino sugar metabolism and peptidoglycan biosynthesis (both *q* < 0.001) **(Fig. 5D)**. These results imply that CST1A and CST1B showed differences in taxonomic composition and metabolic capabilities, supporting their ecological complementarity within the gastric environment.

**Figure 5.**
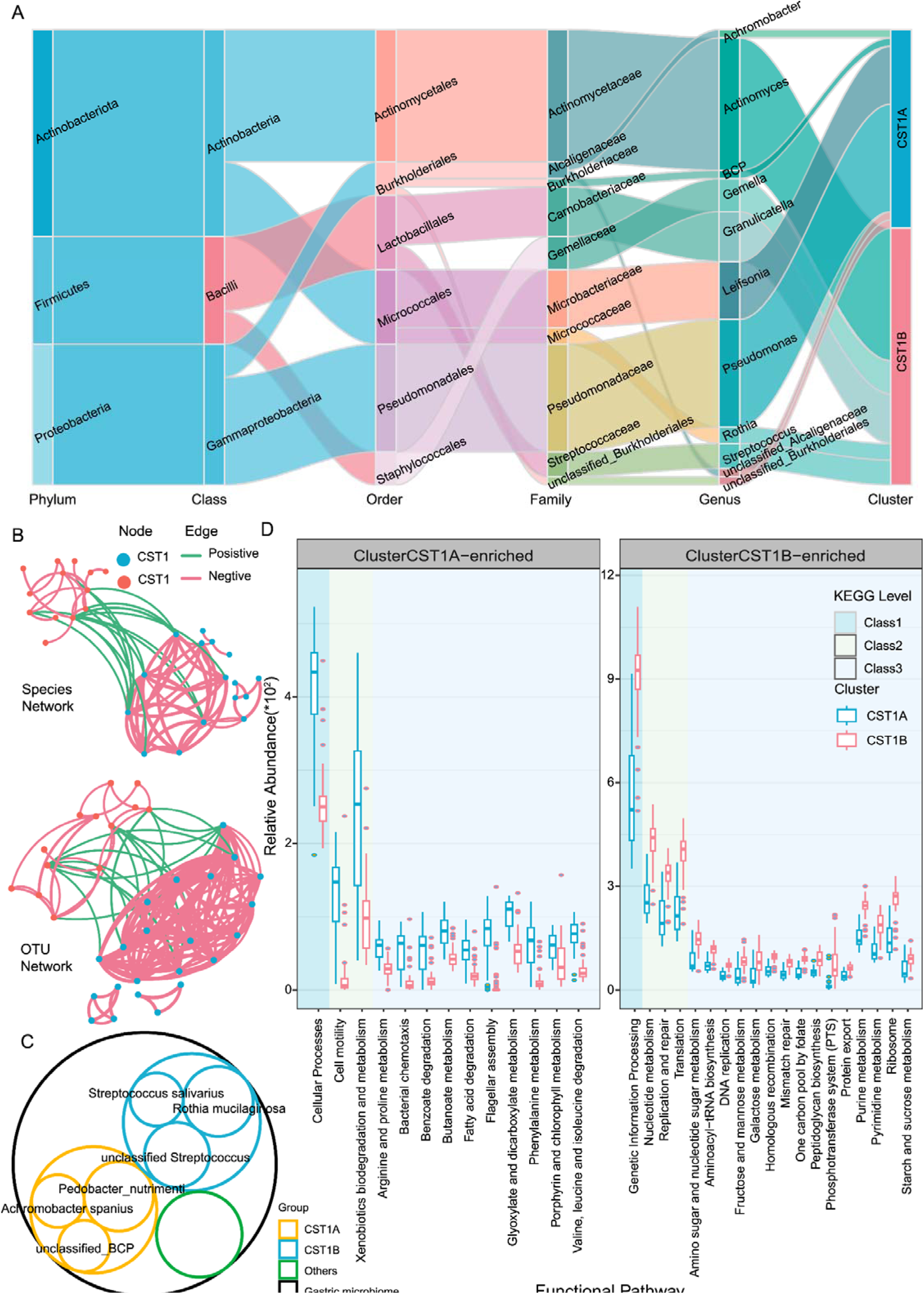
Phylogenetic distribution and functional divergence of core microbial modules. (A) Sankey diagram showing the taxonomic flow of core genera from phylum to genus level, culminating in CST1A and CST1B module assignments. The diagram visualizes hierarchical affiliations across five taxonomic ranks (phylum, class, order, family, genus), highlighting the distinct lineage compositions of each module. (B) Multi-level co-occurrence networks were constructed at the genus, species, and OTU levels based on Spearman correlation (r > 0.5, q < 0.05). Intra-module correlations (within CST1A or CST1B) were consistently positive, while inter-module correlations were predominantly negative, indicating a preserved antagonistic modular structure. (C) Circular stacked chart showing the relative species-level composition of each module. CST1A was dominated by *Pedobacter nutrimenti* (14%), *Achromobacter spanius* (8%), and *unclassified Burkholderia Caballeronia Paraburkholderia* (*BCP*) (7%); CST1B was dominated by *Rothia mucilaginosa* (14%), *unclassified Streptococcus* (12%), and *Streptococcus salivarius* (8%). (D) Faceted box plots displaying the relative abundance of KEGG functional pathways across CST1A and CST1B modules. Left panels show CST1A-enriched pathways (e.g., cell motility, fatty acid degradation), and right panels show CST1B-enriched pathways (e.g., amino sugar metabolism, peptidoglycan biosynthesis). Background shading denotes KEGG pathway categories; box colors distinguish module groupings. Y-axis represents relative pathway abundance; X-axis denotes KEGG pathway names. Statistical comparisons are based on Wilcoxon rank-sum test with Benjamini-Hochberg correction. Comparisons between subgroups were performed using the Wilcoxon rank-sum test. Significance levels are denoted by asterisks: \**q* < 0.05, \*\**q* < 0.01, \*\*\**q* < 0.001 (Benjamini-Hochberg multiple testing).

### Cross-cohort validation reveals structural and predictive conservation of the core gastric microbiota

To evaluate the conservation of the core gastric microbial network across age groups and disease spectrums, we validated this in both a pediatric cohort with duodenal ulcer (n = 51) and five adult cohort with gastric cancer (n = 1,132) **(Fig. 6A)**. For the six cohorts, genus-level abundance profiles of the previously defined core network members were extracted and used as input features for XGBoost classifiers **(Fig. 6B)**. The models demonstrated consistently high predictive performance, with AUC values ranging from 0.694 to 0.944 **(Fig. 6C)**. These results underscore the wide applicability and diagnostic potential of the core microbial network across age, geography, and disease progression. To control for feature selection bias, we constructed comparison models using randomly selected top-N abundant genera (“other” group) and *H. pylori* abundance alone (“Hp” group) for each dataset. ROC analyses showed that the core-based classifier consistently outperformed both alternatives **(Fig. S2F)**. The selected features demonstrated consistently favorable AUC values across various machine learning models, highlighting their robust predictive performance **(Fig. S2G)**. In parallel, these antagonistic network patterns remained consistent across all six independent cohorts: the CST1B subgroup demonstrated robust intra-subgroup positive correlations, whereas CST1A-CST1B inter-subgroup correlations were predominantly negative. This preserved topological architecture collectively supports their role as an evolutionarily maintained homeostatic regulatory module **(Fig. 6D)**. Together, these findings confirm that the gastric core microbial network exhibits both structural stability and predictive generalizability across independent cohorts, highlighting its potential as a robust framework for gastric disease stratification.

**Figure 6.**
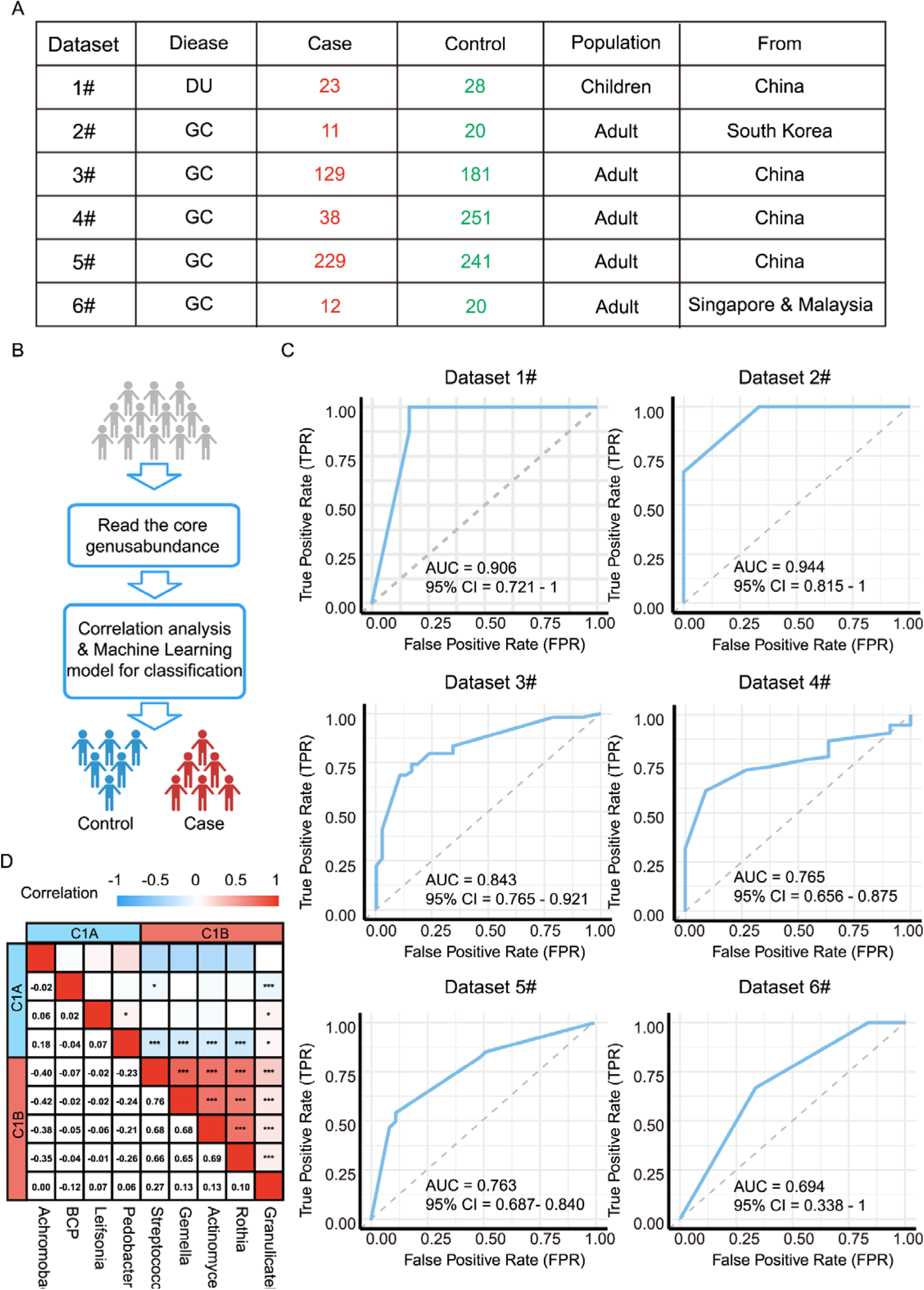
Cross-cohort validation of the ecological stability of the gastric core microbial network. (A) Summary table of six independent gastric microbiome cohorts (n = 1183), including one pediatric duodenal ulcer cohort (Dataset 6, n = 51) and five adult cohorts covering gastric cancer and precancerous lesions (n = 1132). Table columns include dataset ID, disease type, number of cases and controls, population characteristics, and geographical origin. (B) Schematic workflow of the cross-cohort validation strategy. Core network genera identified in the discovery cohort were used to construct genus-level abundance matrices for each validation dataset. XGBoost models were trained in each cohort to classify cases versus controls based on core genera features. (C) Receiver operating characteristic (ROC) curves of XGBoost classifiers across the six cohorts. Area under the curve (AUC) values ranged from 0.724 to 0.896, demonstrating stable classification performance across cohorts with distinct population backgrounds and sequencing platforms. (D) Correlation heatmap showing pairwise Spearman correlations among the 11 core genera based on combined data from all six cohorts. Positive correlations were observed within modules (CST1A and CST1B), while inter-module correlations were predominantly negative, confirming the preserved antagonistic modular structure of the core network across populations. Color scale represents Spearman correlation coefficients ranging from -1 (blue) to +1 (red). Significance levels are denoted by asterisks: \**q* < 0.05, \*\**q* < 0.01, \*\*\**q* < 0.001 (Benjamini-Hochberg multiple testing).

## Discussion

In this study, we systematically constructed and validated a core gastric microbiota network that is established during childhood and preserved across age groups and disease stages. Utilizing full-length 16S/ITS amplicon sequencing and ecological co-occurrence modeling of 246 pediatric gastric biopsy samples, we identified a “seesaw-like” modular network comprising two antagonistic and functionally divergent bacterial clusters, termed CST1A and CST1B. This network structure, consistently detectable across superficial gastritis, duodenal ulcer, and *H. pylori* infection, represents a putative ecological backbone that maintains gastric homeostasis.

CST1A was enriched with *Burkholderia-Caballeronia-Paraburkholderia* (*BCP*), *Achromobacter*, and *Leifsonia*. *BCP* has been negatively associated with BMI during early intestinal development^22^ and is implicated in bile acid metabolism^23^. *Achromobacter* is depleted in patients with cirrhosis, suggesting its potential role as a marker of mucosal integrity^24^. *Leifsonia* commonly found on healthy skin and oral mucosa, may possess ecological migration capacity contributing to gastric mucosal repair^25,26^. In terms of functionality, CST1A is characterized by the enrichment of fatty acid degradation pathways and bacterial motility functions, suggesting strong environmental adaptability and metabolic activity that may contribute to its colonization and ecological stabilization within the gastric mucosa^27,28^. In addition, the core intestinal microbiota is enriched with enzymes that degrade plant polysaccharides and genes essential for butyrate synthesis. These microbial components collectively maintain mucosal homeostasis by producing butyrate and acetate, suppressing inflammatory responses, and enhancing host metabolic functions^14^. These findings show that the core gastric microbiota differs functionally from the intestinal microbiota, suggesting that it operates through distinct protective and pathogenic mechanisms unique to the stomach environment.

CST1B was enriched with oral-origin opportunistic pathogens such as *Streptococcus*, *Rothia*, and *Granulicatella*, whose predicted functions associated with proliferation and adhesion. Prior studies suggest that these genera synergize with *H. pylori* to activate inflammatory MAPK signaling and disrupt the cell cycle, facilitating preneoplastic transformation^29^. These taxonomic and functional features collectively situate CST1B within a broader ecological context that mirrors the intestinal pathobiont guild, characterized by elevated metabolic demands and ecological disruptiveness. The pronounced enrichment of ribosomal components, transporter systems, and nucleic acid metabolism pathways further suggests high biosynthetic and proliferative capacity, which may contribute to ecological disruption under conditions of nutrient competition or microbial imbalance. CST1B may therefore be regarded as a metabolically demanding, ecologically disruptive cluster within the gastric microbiota.

Similar with adult gastric microbiota profiles, taxa linked to the CST1B module are predominantly oral-derived, as evidenced by the prevalence of *Streptococcus*, *Neisseria*, and *Fusobacterium* in prior studies. Oral-derived bacteria have been shown to be persistently enriched in gastric cancer patients^4,30,31^, with *Streptococcus anginosus* in particular being closely associated with histopathological progression^9^. The longitudinal eradication studies also have linked sustained colonization by oral taxa to the emergence and persistence of precancerous lesions such as atrophic gastritis and intestinal metaplasia, suggesting a functional role for these microbes in early tumorigenic transitions^30^. Notably, the pediatric focused approach captures the early-life assembly of the gastric microbiota under near physiological conditions, minimizing confounders and enabling the identification of previously overlooked protective modules. Beyond the well-recognized enrichment of gastric pathogens, we identified and characterized a previously underappreciated microbial module (CST1A), whose members remain ecologically stable in the pediatric stomach yet become markedly depleted during the acute phase of duodenal ulcer.

The XGBoost model trained on core genera achieved robust classification performance across the cohorts (AUC = 0.694-0.944). We propose ecological hypothesis wherein CST1A functions as a foundational module that safeguards mucosal equilibrium and is selectively vulnerable to pathogenic disruption. Its depletion may signal the early onset of gastric disease trajectories. As such, CST1A represents not only a potential biomarker for risk stratification but also a mechanistically actionable target for microbiota-based interventions. This highlights the broader relevance of core microbial networks as evolutionarily preserved scaffolds that underpin the transition from physiological stability to ecological disruption across host developmental stages.

In summary, we present a structurally coherent, functionally antagonistic, and disease preserved core gastric microbiota network that orchestrates ecological balance and is perturbed in the course of gastric disease. This network provides a new framework to understand the progression from *H. pylori* colonization to disease via “single pathogen to network imbalance” transition and offers a theoretical and practical basis for microbiota informed risk prediction and precision intervention strategies. While this study delineates key ecological dynamics of core gastric microbiota, the limitations warrant acknowledgment. First, the mechanistic basis of microbe-host cell interactions remains unexplored, and future studies employing spatially resolved multi-omics approaches (e.g., single cell RNA-seq coupled with microbiome mapping) should prioritize elucidating these crosstalk networks. Second, the absence of interventional cohorts with serial sampling precludes causal inference, representing a critical gap that longitudinal trials would address by systematically tracking microbiota disease temporal causality.

## Materials and methods

### Sample collection and DNA preparation

A total of 246 pediatric participants were recruited from Fujian Children’s Hospital, including 187 children with no endoscopic signs of gastric disease and 59 diagnosed with duodenal ulcers. Gastric mucosal samples were collected using sterile cytology brushes during upper endoscopy. The brush head was aseptically cut with autoclaved scissors, placed into sterile cryovials, snap-frozen in liquid nitrogen, and transferred to -80 °C within 1 hour for long-term storage. The study protocol was approved by the Ethics Committee of Fujian Medical University (Approval No. 2024-232) and Fujian Provincial Children’s Hospital (Approval No. 2024ETKLRK12011). All participants provided written informed consent.

Samples were mechanically lysed using zirconia bead-beating, followed by enzymatic digestion with lysozyme. Total genomic DNA was extracted using the Magnetic Universal Genomic DNA Kit (TIANGEN, China) according to the manufacturer’s instructions. DNA concentration and purity were assessed with a NanoDrop 2000 spectrophotometer, and only samples with OD_260/280_ values between 1.8 and 2.0 were retained. Full-length 16S rRNA and ITS amplicons were amplified using the Kinnex 16S rRNA Kit and Kinnex PCR 12-fold Kit, respectively. Primer sequences (with barcodes) were as follows:

16S Forward:

CTACACGACGCTCTTCCGATCTGATCGAGTCAAGRGTTYGATYM TGGCTCAG

16S Reverse:

AAGCAGTGGTATCAACGCAGAGTCATCGACGTRGYTACCTTGTT ACGACTT

ITS Forward:

CTACACGACGCTCTTCCGATCTTATCGGTGCACTTGGTCATTTAGAGGAAGTAA

ITS Reverse:

AAGCAGTGGTATCAACGCAGAGCTGCGTAACTCCGTGTTTCAAGACGGG

### Publicly Available Datasets

We collected six publicly available 16S rRNA sequencing datasets from gastric samples, designated as Dataset 1#^32^, Dataset 2#^33^, Dataset 3#^34^, Dataset 4#^4^, Dataset 5#^2^, and Dataset 6#^35^. All datasets applied uniform exclusion criteria: no antibiotic, proton pump inhibitor, or probiotic use within 1 month prior to sampling. Raw FASTQ files were downloaded from the SRA database and processed through a unified analysis pipeline.

#### Sequence Processing and Taxonomic Annotation

HiFi reads were generated using the PacBio Revio platform. Sequence processing was performed using QIIME2 (v2020.11)^36^. Reads were merged, quality-filtered, and dereplicated using VSEARCH and Deblur plugins to construct feature tables and representative sequences. Taxonomic classification was conducted using a QIIME2-integrated classifier trained on the SILVA 138 reference database.

#### Statistical Analysis

To evaluate group-level differences while controlling for potential confounders, multivariate linear regression models were applied to compare superficial gastritis controls and duodenal ulcer patients. Hematological parameters were treated as dependent variables, with disease status as the primary predictor, adjusting for age, sex, and delivery mode (vaginal vs. cesarean). Regression coefficients and 95% confidence intervals were reported. Statistical significance was defined as a two-sided q-value < 0.05. All analyses were conducted using R version 4.2.1.

Microbial abundance data were analyzed using MaAsLin2, incorporating age, sex, and delivery mode as covariates. Bacterial genera were considered significantly different if |β| > 0.1 and FDR < 0.05.

#### Microbial Association Network Analysis

At the genus level, co-occurrence networks were constructed based on pairwise Spearman correlations (|ρ| ≥ 0.5, FDR < 0.05). Genera pairs showing significant positive or negative correlations were included. Networks were visualized in Gephi (v0.9.2), where nodes represented genera and edges represented significant correlations, with edge weight proportional to the correlation coefficient.

#### Microbial Community Typing

Microbial community types were identified using Partitioning Around Medoids (PAM) clustering based on Jensen-Shannon divergence and Bray-Curtis dissimilarity matrices^37^. Clustering was performed using the cluster and factoextra R packages, and the optimal number of clusters was determined by silhouette width. Cluster robustness was validated by k-means clustering and Rand index analysis. Representative genera from each cluster were used in downstream functional and clinical association analyses.

#### Prediction of Microbial Functional Potential

Functional profiles of the 16S data were inferred using PICRUSt2, with annotations mapped to KEGG Orthology (KO) terms. Functional abundances were grouped by microbial cluster, and inter-group comparisons were conducted using the Wilcoxon rank-sum test. Pathways with FDR < 0.05 were considered significantly different.

#### Machine Learning-Based Classification

To evaluate the classification performance of core microbiota features, we constructed seven machine learning models, including XGBoost, Random Forest, AdaBoost, Neural Network, GBM, and GBMBoost. All models were trained using a 7:3 train-test split, and input features were standardized prior to modeling.

Model-specific parameters were configured accordingly, and training was performed on the training set. For the XGBoost model, 10-fold cross-validation was applied to determine the optimal number of boosting rounds and to prevent overfitting. Model performance was evaluated on the independent test set using receiver operating characteristic (ROC) curves and the area under the curve (AUC), with 95% confidence intervals calculated via bootstrapping. In addition, we performed comparative modeling using feature sets excluding *Helicobacter* or containing only *Helicobacter* to assess the independent predictive capacity of the core microbiota features across all algorithms.

## Data and software availability

The amplicon sequencing data generated in this study have been deposited in the NCBI Sequence Read Archive (SRA) under BioProject accession numbers PRJNA1251565 (16S rRNA amplicon data) and PRJNA1251792 (ITS amplicon data).

## Author contributions

Z.W.X., L.X and H.W.J. conceived the idea. W.D.H. performed nucleic acid extraction of microbiota, quality control, and data analysis. H.W.J. conducted microbial community sequencing and quality control procedures. X.J.G., Y.Y.S., H.Z., J.Y.S., T.Z., Y.B.C., Y.Y.Y. and W.W. conducted clinical sample collection and quality control. Z.W.X., L.X, H.W.J. and W.D.H. prepared the figures. Z.W.X. and W.D.H. wrote the manuscript.

## Disclosure and competing interest statement

The authors declare no competing interests.

## Acknowledgements

This study was supported by the Fujian Province Key Technological Innovation Research and Industrialization Project, China (No. 2024XQ012), the Natural Science Foundation of Fujian Province, China (No. 2022J01197).

---

**Figure S1. Quality control and preprocessing of clinical and microbial data.**

(A) Comparison of *Helicobacter pylori* detection results across four methods: rapid urease test (RUT), immunohistochemistry (IHC), ¹³ C-urea breath test (C13), and third-generation sequencing (Seq). The heatmap presents detection outcomes across all samples (columns), with red indicating positive results. ROC analysis based on sequencing-derived relative abundance yielded an optimal diagnostic threshold of 0.01 for *H. pylori* positivity, achieving an AUC of 0.927 (95% CI: 0.855-1.000).

(B) Boxplots comparing hematological parameters between duodenal ulcer (DU) and superficial gastritis (SG) patients through multiple linear regression models adjusted for age, sex, and birth mode. DU patients exhibited significantly lower mean corpuscular hemoglobin concentration (MCHC, q<0.05), hematocrit (HCT, q<0.05), red blood cell count (RBC, q<0.01), and hemoglobin concentration (HGB, q<0.001), while showing increased immature granulocyte count (IG count, p<0.05), red cell distribution width (RDW, q<0.01), and RDW coefficient of variation (q<0.01).

(C) Threshold determination for bacterial data denoising. Based on retention curves, a relative abundance threshold of 0.005 was selected (species with abundance <0.005 set to zero), and a prevalence threshold of 0.01 was applied (species retained if present in ≥10% of samples).

(D) Threshold determination for fungal data denoising. Using a similar strategy, the abundance threshold was set at 0.006 and the prevalence threshold at 0.08 (species retained if present in ≥8% of samples).

**Figure S2. Stratification of *H. pylori*-negative controls and machine learning-based classification in *H. pylori*-positive individuals.**

(A) PAM clustering based on the abundance profiles of 11 previously defined core genera identified two optimal subgroups (K = 2) within the *H. pylori*-negative non-ulcer control (HpNNC) cohort, as indicated by silhouette width analysis.

(B) Validation of clustering robustness using k-means algorithm, showing high concordance with PAM clustering (Adjusted Rand Index, ARI = 0.84).

(C) Panel of four aligned plots comparing HpNNC1 and HpPNC groups using core genera as variables: (1) bar plot of model-derived coefficients (effect size) for each genus; (2-3) genus-level prevalence in HpNNC1 and HpPNC groups; (4) boxplots showing relative abundance distributions, highlighting shifts in community structure. Microbial abundance differential analysis was performed using MaAsLin2(v1.6.0) with robust linear models with fixed effects comprising disease status, age, gender, and birth mode. Raw abundances were normalized by sum scaling, with significance thresholds defined as log2(fold change)>1 and FDR-adjusted q<0.05. Differential taxa are annotated in figures as: * *q*<0.05, \**q*<0.01, *** *q*<0.001 (Benjamini-Hochberg correction).

(D) ROC curve of XGBoost model trained on core genera to distinguish between *H. pylori*-positive non-ulcer (HpPNC) and duodenal ulcer (HpPDU) groups, achieving excellent performance (AUC = 0.937, 95% CI: 0.886-0.987).

(E) Feature importance plot showing contributions of individual core genera to the classifier’s performance, supporting their discriminatory value despite the absence of statistically significant differential taxa.

(F) Paired boxplots comparing model predictive performance among the Core feature group (core), the Bias-excluded group (other), and the *Helicobacter pylori* group (Hp). Boxes represent the interquartile range (IQR), horizontal lines indicate medians, whiskers extend to 1.5×IQR, and discrete points denote outliers. Wilcoxon signed-rank tests revealed a statistically significant AUC difference between the Core and Hp groups. (G)Heatmap showing AUC performance of six machine learning models across different datasets. Rows represent classification algorithms, including XGBoost (xgb), Random Forest (rf), AdaBoost (ada), Neural Network (nn), GBM (gbm), and GBMBoost (gbmboost); columns represent individual datasets. Numeric values within the heatmap indicate the AUC achieved by each model on each dataset, highlighting the robustness and consistency of model performance across varying data inputs.

## Supplementary Material

Supplementary Material, page 1: Top 20 Fungal Genera and Relative Abundance in the HpNNC Group.

Supplementary Material, page 2: Spearman Correlation Matrix of Fungal Relative Abundance in the HpPDU Group.

Supplementary Material, pages 3-4: Edge List for Fungal Interaction Networks, including Edge Weight and Correlation Coefficients, Across the HpNNC, HpPNC, and HpNDU Groups.

Supplementary Material, page 5: Core Bacterial Genera Abundance Shows No Significant Variation Across Sex or Delivery Mode.

Supplementary Material, page 6: Comparative Analysis of Fungal Community Composition Across Groups with No Significant Differences

## Notes

### Competing Interest Statement

The authors have declared no competing interest.

