## Supplementary figures and images for "Core gastric microbiota linked to pathogenesis and preserved across age stratified cohorts"

### Supplemental Fig.1

#### Slide 1
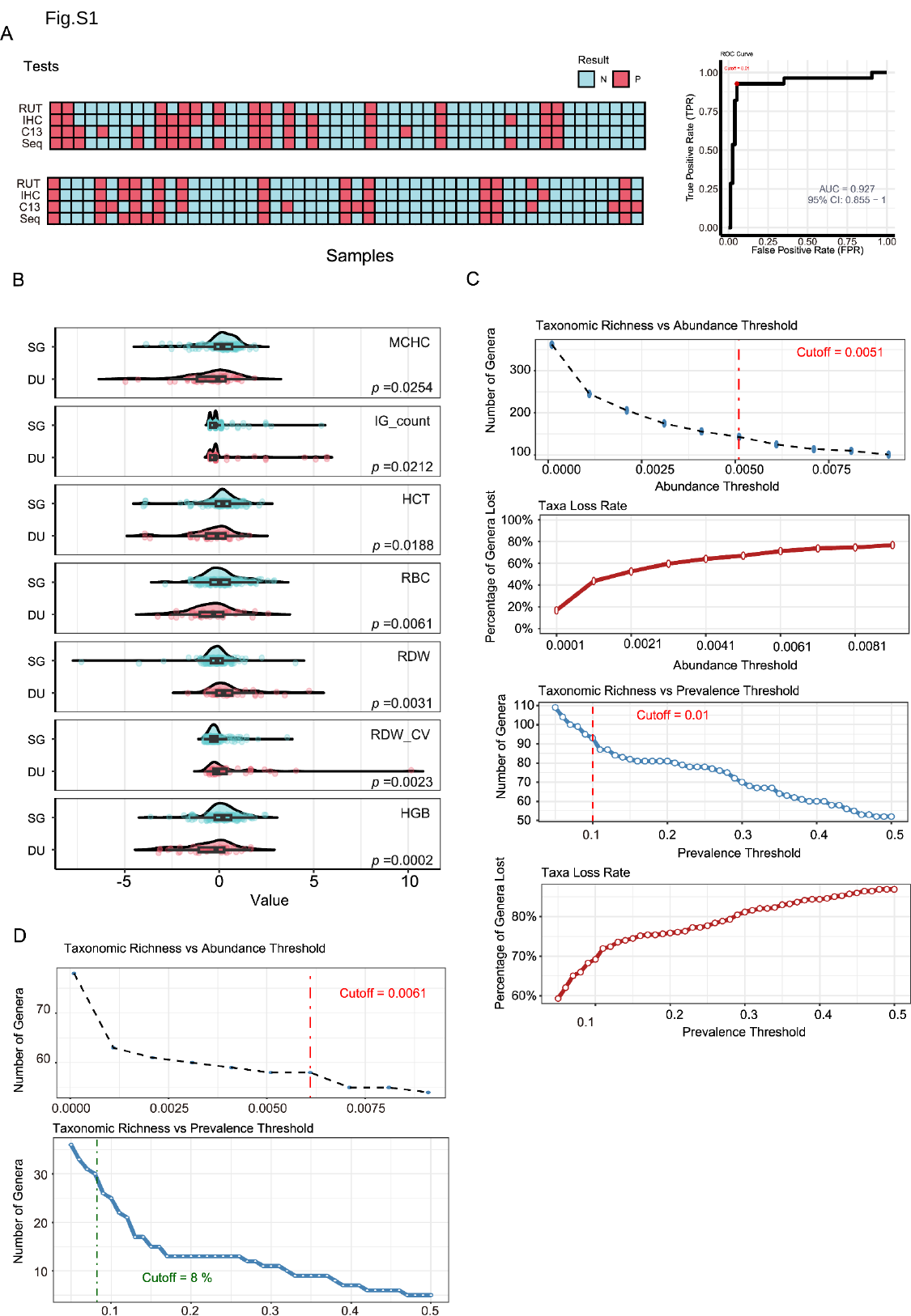

Fig.S1

#### Slide 2
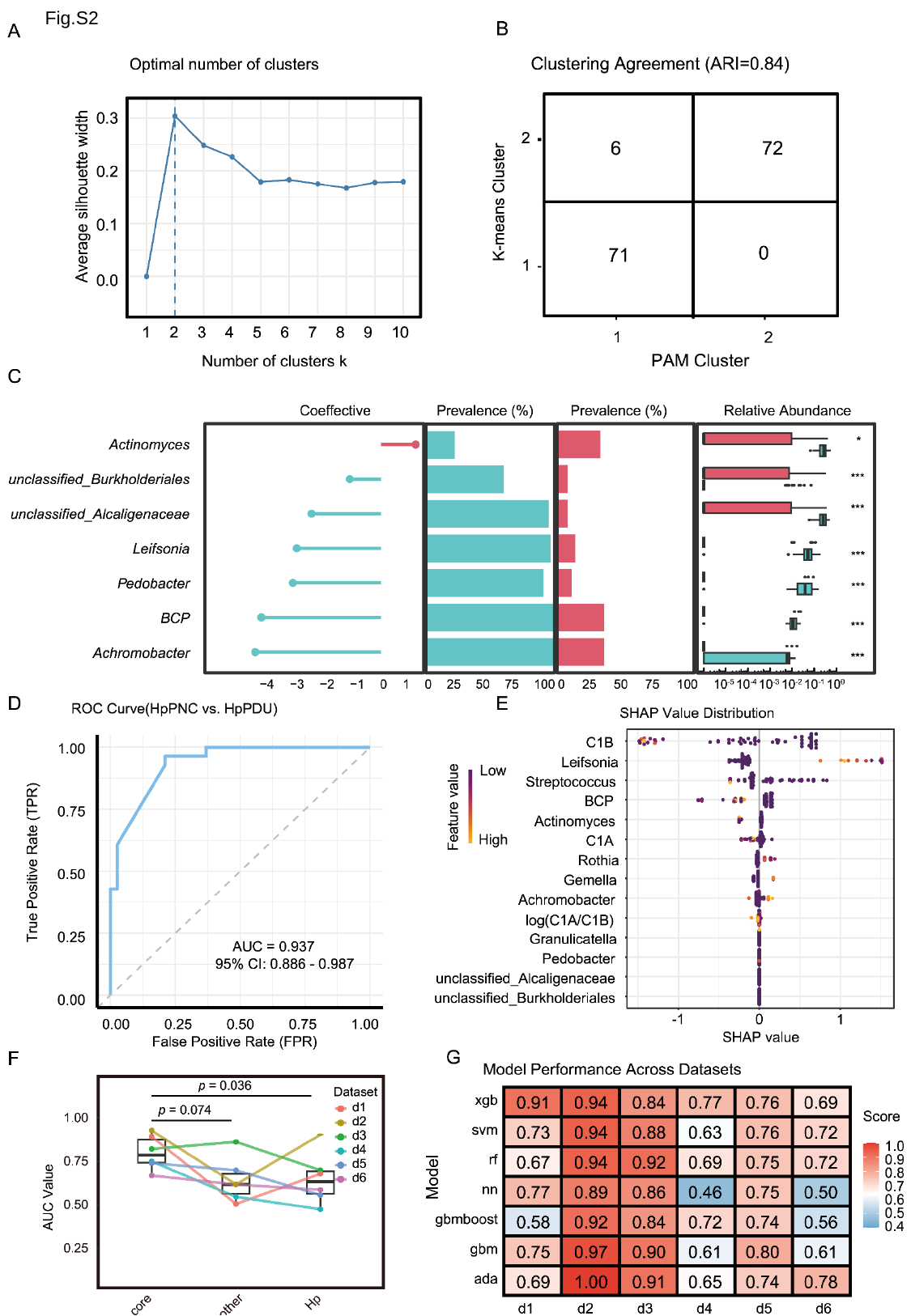

Fig.S2
