## Supplemental Material for "Core gastric microbiota linked to pathogenesis and preserved across age stratified cohorts"

**Supplementary Material**

**Supplementary Material, page 1:** Top 20 Fungal Genera and Relative Abundance in the HpNNC Group.

**Supplementary Material, page 2:** Spearman Correlation Matrix of Fungal Relative Abundance in the HpPDU Group.

**Supplementary Material, pages 3-4:** Edge List for Fungal Interaction Networks, including Edge Weight and Correlation Coefficients, Across the HpNNC, HpPNC, and HpNDU Groups.

**Supplementary Material, page 5:** Core Bacterial Genera Abundance Shows No Significant Variation Across Sex or Delivery Mode.

**Supplementary Material, page 6:** Comparative Analysis of Fungal Community Composition Across Groups with No Significant Differences.

| No. | Genus | AverageAbundance |
| --- | --- | --- |
| 1 | Malassezia | 0.3810 |
| 2 | Candida | 0.1486 |
| 3 | Cutaneotrichosporon | 0.0547 |
| 4 | Aspergillus | 0.0400 |
| 5 | Trametes | 0.0344 |
| 6 | Setophaeosphaeria | 0.0267 |
| 7 | Archaeorhizomyces | 0.0252 |
| 8 | Fungi_gen_Incertae_sedis | 0.0225 |
| 9 | Hygrophorus | 0.0225 |
| 10 | Cladosporium | 0.0200 |
| 11 | Rhodotorula | 0.0198 |
| 12 | Cryptococcus | 0.0137 |
| 13 | Sordariomycetes_gen_Incertae_sedis | 0.0121 |
| 14 | unclassified_Fungi | 0.0118 |
| 15 | Russula | 0.0117 |
| 16 | Malasseziaceae_gen_Incertae_sedis | 0.0110 |
| 17 | Aureobasidium | 0.0106 |
| 18 | Hypocreales_gen_Incertae_sedis | 0.0103 |
| 19 | Peniophorella | 0.0092 |
| 20 | unclassified_Chytridiomycota | 0.0092 |

Supplementary Material, page 1 : Top 20 Fungal Genera and Relative Abundance in the HpNNC Group.

|  | Aspergillus | Candida | Fungi_gen_Incertae_sedis | Malassezia | Malasseziaceae_gen_Incertae_sedis | Neurospora | Wallemia | unclassified_Fungi |
| --- | --- | --- | --- | --- | --- | --- | --- | --- |
| Aspergillus | 0 | 0 | 0 | 0 | 0 | 0 | 0 | 0 |
| Candida | 0 | 0 | 0 | 0 | 0 | 0 | 0 | 0 |
| Fungi_gen_Incertae_sedis | 0 | 0 | 0 | 0 | 0 | 0 | 0 | 0 |
| Malassezia | 0 | 0 | 0 | 0 | 0 | 0 | 0 | 0 |
| Malasseziaceae_gen_Incertae_sedis | 0 | 0 | 0 | 0 | 0 | 0 | 0 | 0 |
| Neurospora | 0 | 0 | 0 | 0 | 0 | 0 | 0 | 0 |
| Wallemia | 0 | 0 | 0 | 0 | 0 | 0 | 0 | 0 |
| unclassified_Fungi | 0 | 0 | 0 | 0 | 0 | 0 | 0 | 0 |

Supplementary Material, page 2: Spearman Correlation Matrix of Fungal Relative Abundance in the HpPDU Group.

| Group | source | target | weight | correlation |
| --- | --- | --- | --- | --- |
| HpNNC | Candida | Malassezia | 0.3498 | -0.3498 |
| HpNNC | Trametes | unclassified_Fungi | 0.3437 | 0.3437 |
| HpNDU | Acaulium | Chaetomella | 1.0000 | 1.0000 |
| HpNDU | Acaulium | Fusarium | 1.0000 | 1.0000 |
| HpNDU | Acaulium | Humicola | 1.0000 | 1.0000 |
| HpNDU | Acaulium | Hypocreales_gen_Incertae_sedis | 0.6573 | 0.6573 |
| HpNDU | Acaulium | Staphylotrichum | 1.0000 | 1.0000 |
| HpNDU | Acaulium | Trichoderma | 1.0000 | 1.0000 |
| HpNDU | Acaulium | unclassified_Fungi | 0.7230 | 0.7230 |
| HpNDU | Chaetomella | Fusarium | 1.0000 | 1.0000 |
| HpNDU | Chaetomella | Humicola | 1.0000 | 1.0000 |
| HpNDU | Chaetomella | Hypocreales_gen_Incertae_sedis | 0.6573 | 0.6573 |
| HpNDU | Chaetomella | Staphylotrichum | 1.0000 | 1.0000 |
| HpNDU | Chaetomella | Trichoderma | 1.0000 | 1.0000 |
| HpNDU | Chaetomella | unclassified_Fungi | 0.7230 | 0.7230 |
| HpNDU | Cladosporium | Neurospora | 1.0000 | 1.0000 |
| HpNDU | Cutaneotrichosporon | unclassified_Fungi | 0.6573 | 0.6573 |
| HpNDU | Fusarium | Humicola | 1.0000 | 1.0000 |
| HpNDU | Fusarium | Hypocreales_gen_Incertae_sedis | 0.6573 | 0.6573 |
| HpNDU | Fusarium | Staphylotrichum | 1.0000 | 1.0000 |
| HpNDU | Fusarium | Trichoderma | 1.0000 | 1.0000 |
| HpNDU | Fusarium | unclassified_Fungi | 0.7230 | 0.7230 |
| HpNDU | Humicola | Hypocreales_gen_Incertae_sedis | 0.6573 | 0.6573 |
| HpNDU | Humicola | Staphylotrichum | 1.0000 | 1.0000 |
| HpNDU | Humicola | Trichoderma | 1.0000 | 1.0000 |
| HpNDU | Humicola | unclassified_Fungi | 0.7230 | 0.7230 |
| HpNDU | Hypocreales_gen_Incertae_sedis | Staphylotrichum | 0.6573 | 0.6573 |
| HpNDU | Hypocreales_gen_Incertae_sedis | Trichoderma | 0.6573 | 0.6573 |
| HpNDU | Meyerozyma | Penicillium | 1.0000 | 1.0000 |
| HpNDU | Meyerozyma | Russula | 1.0000 | 1.0000 |
| HpNDU | Penicillium | Russula | 1.0000 | 1.0000 |
| HpNDU | Schizophyllum | Setophaeosphaeria | 0.6573 | 0.6573 |
| HpNDU | Staphylotrichum | Trichoderma | 1.0000 | 1.0000 |
| HpNDU | Staphylotrichum | unclassified_Fungi | 0.7230 | 0.7230 |
| HpNDU | Trichoderma | unclassified_Fungi | 0.7230 | 0.7230 |
| HpPNC | Archaeorhizomyces | Penicillium | 0.6595 | 0.6595 |
| HpPNC | Articulospora | Tulasnella | 0.7223 | 0.7223 |
| HpPNC | Candida | Penicillium | 0.6595 | 0.6595 |
| HpPNC | Coniothyrium | Saccharomyces | 1.0000 | 1.0000 |
| HpPNC | Lactifluus | Malasseziaceae_gen_Incertae_sedis | 1.0000 | 1.0000 |
| HpPNC | Trametes | Tulasnella | 0.7586 | 0.7586 |

Supplementary Material, page 3-4: Edge List for Fungal Interaction Networks, including Edge Weight and Correlation Coefficients, Across the HpNNC, HpPNC, and HpNDU Groups.

| Variable | Group | Comparison | P_value | Effect_size | Method | Significance |
| --- | --- | --- | --- | --- | --- | --- |
| cluster1 | Gender | Male vs Female | 0.505258298 | 0.05476451 | Mann-Whitney U test | ns |
| cluster2 | Gender | Male vs Female | 0.651679433 | 0.03710734 | Mann-Whitney U test | ns |
| log_ratio | Gender | Male vs Female | 0.391298491 | 0.07046674 | Mann-Whitney U test | ns |
| cluster1 | Birth_mode | CS vs NC | 0.322724499 | 0.081561 | Mann-Whitney U test | ns |
| cluster2 | Birth_mode | CS vs NC | 0.464385882 | 0.06034498 | Mann-Whitney U test | ns |
| log_ratio | Birth_mode | CS vs NC | 0.42141227 | 0.06631062 | Mann-Whitney U test | ns |

Supplementary Material, page 5: Core Bacterial Genera Abundance Shows No Variation Across Sex or Delivery Mode

| feature | Case | Control | coef | stderr | N | N.not.0 | pval | qval |
| --- | --- | --- | --- | --- | --- | --- | --- | --- |
| Trametes | HpNDU | HpNNC1 | -0.232030028 | 0.119056451 | 71 | 8 | 0.055559022 | 0.730329752 |
| Malassezia | HpNDU | HpNNC1 | 0.344269712 | 0.321448977 | 71 | 44 | 0.288073344 | 0.823066696 |
| Aspergillus | HpNDU | HpNNC1 | 0.330215371 | 0.62517703 | 71 | 15 | 0.599135575 | 0.907408966 |
| Candida | HpNDU | HpNNC1 | -0.231735406 | 0.282518194 | 71 | 10 | 0.415026511 | 0.907408966 |
| unclassified_Fungi | HpNDU | HpNNC1 | -0.150839358 | 0.385134975 | 71 | 11 | 0.696576158 | 0.907408966 |
| Candida | HpPNC | HpNNC1 | -0.210539963 | 0.273240059 | 72 | 10 | 0.443693542 | 0.904302554 |
| unclassified_Fungi | HpPNC | HpNNC1 | 0.76389169 | 0.497785773 | 72 | 16 | 0.12959622 | 0.904302554 |
| Aspergillus | HpPNC | HpNNC1 | -0.230590263 | 0.553037955 | 72 | 12 | 0.678046459 | 0.971093822 |
| Malassezia | HpPNC | HpNNC1 | -0.139183365 | 0.479010219 | 72 | 42 | 0.772281917 | 0.971093822 |
| Trametes | HpPNC | HpNNC1 | -0.0310231 | 0.148812888 | 72 | 11 | 0.835494063 | 0.971093822 |
| Malasseziaceae_gen_Incertae_sedis | HpPDU | HpNNC1 | 0.475613736 | 0.150345231 | 73 | 8 | 0.002331054 | 0.055945286 |
| Aspergillus | HpPDU | HpNNC1 | 1.5168639 | 0.637274382 | 73 | 20 | 0.02011077 | 0.241329236 |
| Trametes | HpPDU | HpNNC1 | -0.220559612 | 0.115995425 | 73 | 8 | 0.061480817 | 0.3688849 |
| Malassezia | HpPDU | HpNNC1 | 0.423824502 | 0.407181217 | 73 | 48 | 0.301620574 | 0.804321531 |
| Candida | HpPDU | HpNNC1 | -0.058904244 | 0.309459173 | 73 | 12 | 0.849605303 | 0.970567629 |
| unclassified_Fungi | HpPDU | HpNNC1 | 0.074709517 | 0.401483735 | 73 | 12 | 0.852933246 | 0.970567629 |
| Aspergillus | HpPDU | HpPNC | 1.589846826 | 0.748217852 | 49 | 14 | 0.039253225 | 0.211703581 |
| Malasseziaceae_gen_Incertae_sedis | HpPDU | HpPNC | 0.475773809 | 0.206787925 | 49 | 7 | 0.026201677 | 0.211703581 |
| Malassezia | HpPDU | HpPNC | 0.50407037 | 0.571635219 | 49 | 32 | 0.382674625 | 0.803419978 |
| Wallemia | HpPDU | HpPNC | 0.208988729 | 0.224220908 | 49 | 5 | 0.356388762 | 0.803419978 |
| Candida | HpPDU | HpPNC | 0.038900528 | 0.158243207 | 49 | 6 | 0.806958907 | 0.984638471 |
| unclassified_Fungi | HpPDU | HpPNC | -0.142359467 | 0.529707785 | 49 | 10 | 0.789377499 | 0.984638471 |

Supplementary Material, page 6: Comparative Analysis of Fungal Community Composition Across Groups with No Significant Differences.
